# Deep learning model can predict water binding sites on the surface of proteins using limited-resolution data

**DOI:** 10.1101/2020.04.20.050393

**Authors:** Jan Zaucha, Charlotte A. Softley, Michael Sattler, Grzegorz M. Popowicz

## Abstract

The surfaces of proteins are generally hydrophilic but there have been reports of sites that exhibit an exceptionally high affinity for individual water molecules. Not only do such molecules often fulfil critical biological functions, but also, they may alter the binding of newly designed drugs. In crystal structures, sites consistently occupied in each unit cell yield electron density clouds that represent water molecule presence. These are recorded in virtually all high-resolution structures obtained through X-ray diffraction. In this work, we utilized the wealth of data from the RCSB Protein Data Bank to train a residual deep learning model named ‘hotWater’ to identify sites on the surface of proteins that are most likely to bind water, the so-called water hot spots. The model can be used to score existing water molecules from a PDB file to provide their ranking according to the predicted binding strength or to scan the surface of a protein to determine the most likely water hot-spots *de novo*. This is computationally much more efficient than currently used molecular dynamics simulations. Based on testing the model on three example proteins, which have been resolved using both high-resolution X-ray crystallography (providing accurate positions of trapped waters) as well as low-resolution X-ray diffraction, NMR or CryoEM (where structure refinement does not yield water positions), we were able to show that the hotWater method is able to recover in the “water-free” structures many water binding sites known from the high-resolution structures. A blind test on a newly solved protein structure with waters removed from the PDB also showed good prediction of the crystal water positions. This was compared to two known algorithms that use electron density and was shown to have higher recall at resolutions >2.6 Å. We also show that the algorithm can be applied to novel proteins such as the RNA polymerase complex from SARS-CoV-2, which could be of use in drug discovery. The hotWater model is freely available at (https://pypi.org/project/hotWater/).

## 1. Introduction

Water is the “matrix of life”, as referred to by Szent-Győrgi [1]. With regard to proteins, not only does it provide the necessary energetic conditions required for faithful folding [2–4] but also mediates enzymatic activity and facilitates substrate specificity [5,6]. In addition, there are instances of specific water molecules that fulfill critical biological roles contributing to a protein’s conformational stability [7], enabling proton translocation in the membrane-bound transhydrogenase [8], mediating an antigen-antibody association [9], or manifestly contributing to ligand binding [10–12]. Recently, we demonstrated that considering the particularly strongly interacting water molecules when designing drugs allows the identification of ligands that are specific to a particular protein ortholog [13].

Although the surface of most proteins is generally hydrophilic, interactions between water molecules and atoms comprising the protein are transient. Nevertheless, upon crystallization of a protein for the purpose of structure determination, it is evident from the obtained electron density clouds that many water molecules occupy conserved positions within the repetitive crystal lattice [14]. It is a statistical indication that water molecules are bound at these specific positions on the surface of the protein and points to a high affinity of these sites to water. Such water molecules are present in virtually all structures resolved to high resolution by X-ray diffraction [15]. However, water molecules are not typically detectable in low-precision X-ray structures (resolution of more than 3.0Å), and are not present in structures obtained using other experimental techniques such as nuclear magnetic resonance (NMR) and cryogenic electron microscopy (Cryo-EM) [16]. Previous research elucidated the factors affecting the number of water molecules resolved within a crystal [15,17]. These include the crystallographic resolution [15], average temperature factor of the protein’s atoms, the structure’s *R* value, as well as the percentage of solvent in the crystal [17]. In this study, we attempted to go a step further and capture the landscape of water interaction with the protein surface, at the resolution of individual water molecules.

Convolutional neural networks have been repeatedly shown to excel in solving the tasks of image recognition [18–21], speech recognition [22–25] and even genetic or protein sequence analysis [26–33]. In the field of computational chemistry, they have been applied to predict ligand binding affinities [34–37] as well as to identify binding pockets within protein structures[38]. These successes have been attributed to their ability of hierarchically learning attributes indicative of each target class.

With sufficiently large training datasets, increasingly more complex problems can be solved using ever deeper network architectures. However, training very deep networks is often hindered by the “vanishing gradients” phenomenon [39], which arises as a result of ever decreasing partial derivatives of the loss function with respect to the nodes’ weights, preventing further updates and (thus training) of deeply hidden network layers. This has been overcome by the residual network architecture (ResNet) [20,27], which incorporates “skip connectors” that bypass a number of layers and allow the reuse of activations from previous sections of the network that still feature considerable gradient levels. This way, information is efficiently propagated to all parts of the network facilitating the training process.

Encouraged by the success of ResNets in other application domains, we trained such a model, which, given a position near the protein’s surface, provides a score indicating the probability of an interaction with a water molecule. The model can be used to score the existing water molecules within a crystal structure, in order to identify the positions featuring highest affinity for individual water molecules, the so-called ‘water hot-spots’. Furthermore, we demonstrate that the model can be used for scanning the surface of proteins resolved using low resolution X-ray crystallography, NMR or Cryo-EM, to accurately reconstruct the most likely water binding sites. The ability of the model to “decorate” the NMR and Cryo-EM structures with solvent has a variety of applications and does not require time- and computing power-intensive molecular dynamics simulations.

## 2. Materials & Methods

### 2.1 Training dataset

In order to develop a binary regression model, training data providing the positive (empirically supported water binding site) and negative (site with no evidence of binding water) class examples are required.

The positive class data (water molecules from PDB structures) were extracted from a non-redundant representative set (sequence similarity <30%) of 9,067 PDB structures not containing nucleic acids, resolved to at least 1.8Å resolution using X-ray diffraction (using the download tool provided by PDB [40]; data downloaded on 13.04.19; a list of all PDB structures included in the dataset is provided in Supplementary File 1). Water molecules not interacting with at least two atoms (proximity of water’s oxygen to target atoms <4.5Å) of the protein were excluded. Due to computational resource constraints, the positive dataset was limited to 2,800,000 individual water molecules.

Generating the negative class data poses a challenge since the number of positions not occupied by water molecules is virtually infinite. Selecting any positions not occupied by a water molecule would yield a mostly trivial dataset - positions within the core of the protein structure that are physically inaccessible to water molecules or positions too far away from any atoms of the structure. Therefore, the negative class samples were chosen using a heuristic approach, by randomly selecting positions near the protein’s estimated solvation envelope as follows. Firstly, EDTSurf [41] was used to generate a triangular mesh matching the Coulombic radius of water (1.4 Å) around the protein (run with option “–f 20” to use the finest grid available) [42]. Based on empirical evidence, we found that the majority of water molecules in PDB structures can be found within 2.4-5.8Å of the mesh points (this makes sense given that the length of the hydrogen bond is 1.5-4Å [43]). The mesh was scaled outwards and inwards to a depth of 1.8Å to cover cavities within the protein, using increments matching the mean grid size (average distance between two adjacent grid points, typically around 0.2Å) to form multiple mesh layers around the protein (Figure 1). Negative class positions were then randomly sampled from the obtained layers, according to a probability distribution selected to estimate the distribution observed empirically (Gaussian distribution centered at layer corresponding to roughly 2.4Å from the protein’s surface, scaled by a factor of 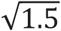 times the number of grid layers). The selected points were verified to check that they were not closer than 1.4Å to a water molecule encoded within the PDB file. The number of negative class samples generated was selected to match the number of positive samples, yielding a balanced training set.

**Figure 1:**
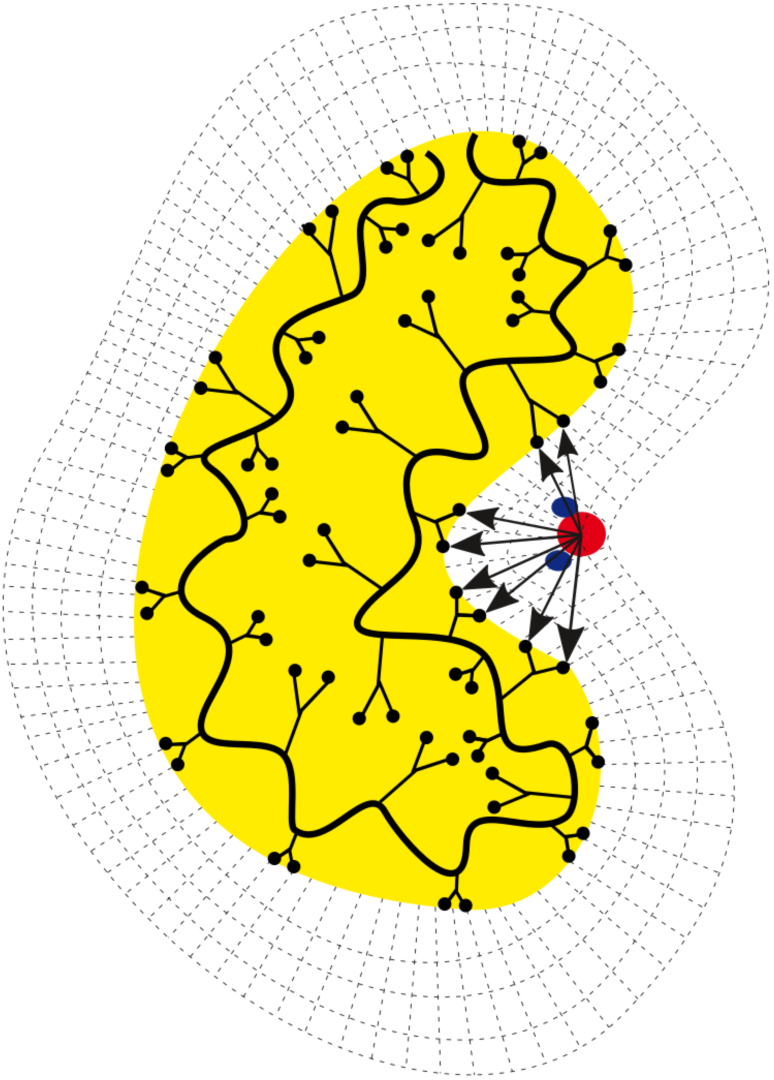
Schematic diagram of mesh layers covering the area around a protein surface. Water molecules present within the grid are encoded in terms of an array of vectors pointing towards all proximal (<=7.5Å) atoms of the protein.

A random subset of samples, amounting to roughly 15% of the training set, was defined as the “holdout” test set and excluded from the training and cross validation steps, for use once the final network architecture and model parameters had been defined. A list of PDB files used for the test set is available in Supplementary File 1.

### 2.2 Input features and data preprocessing

In this work, we have made an arbitrary, but intuitive, decision to encode the queried potential water binding site on the surface of the protein in terms of a linear array of vectors representing the full atomic neighborhood of the site (providing information on the distance to each atom as well as the corresponding atom type and temperature factor). Each positive and negative class sample was annotated with vectors pointing towards all proximal (<=7.5Å) atoms of the protein. The target atom types were recorded according to their chemical properties, distinguishing between: hydrogens, aliphatic carbons, alpha carbons, aromatic carbons, carbonyl carbons of the main chain, amide carbons of the side chain, amide carbons of the main chain, carboxyl carbons, guanidinium carbons, hydroxyl oxygens, carboxyl oxygens, carbonyl oxygens of the main chain, hydroxyl aromatic oxygens, amide nitrogens of the main chain, aromatic nitrogens of tryptophan, aromatic nitrogens of histidine, primary amine nitrogens, secondary amine nitrogens, sulfurs of methionine and sulfurs of cysteine. These 21 atom types are represented in the model using the “one hot encoding”. Since temperature factors (B-values) have been shown to affect the probability of resolving water molecules within the crystal [17], their normalized values were added as additional inputs into the model.

The model was designed to consist of two input channels: the first layer provides vectors pointing from the (non-)water molecule position to the nearest fifty interacting atoms of the protein (shape: 50×3), while the second layer provides B-values and atom-type information corresponding to each of the fifty vectors (shape: 50×30, one hot encoding).

In order to facilitate model convergence and prediction performance, the landscape of input feature patterns that can appear should be condensed; in particular, rotational transformations of the input feature space introduce redundancies which greatly hinder the ability of the model to learn [44]. One way of mitigating the problem is training-data augmentation including random rotations of the each input feature sample [34] or data-preprocessing to ensure rotational invariance of the inputs. In this work, we opted for the latter solution; the vectors were rotated to a common system of coordinates in Euclidean space by Gram-Schmidt orthonormalization using the vector pointing to the closest atom of the protein 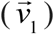 and the orthogonal part of the vector at the highest angle against it 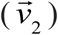 as the first and second basis vectors for the orientation (the third vector forming the new orthonormal basis *ê*_1_, *ê* _2_, *ê* _3_ is their cross product):

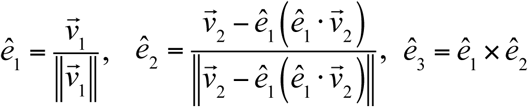

Each vector was rotated into the standardized system of coordinates by left-multiplying it by the matrix *M* = [*ê*_3_, *ê*_2_, *ê* _1_].

In order to allow for effective learning of patterns indicative of strong interactions with the water molecules, the vectors were sorted into ordered pairs such that the first vector is the vector with the lowest norm, the second vector is the vector at the highest angle against it, the next vector pair is again the vector with the lowest norm and the vector at the highest angle against it (drawn from the set of remaining vectors). All data were normalized to fall between [0,1] (B-values were log10-transformed beforehand). If the number of interacting atoms with the water molecule was less than fifty, the unoccupied positions were filled with empty values.

### 2.3 Neural network architecture and hyper-parameter tuning

Due to the vastness of possibilities, the selection of the neural network architecture was inspired by the previous successes in applying similar models to a different application domain [20,27]. The training set was split into six cross-validation folds (excluding the holdout test set) and various network depths (numbers of hidden layers and numbers of residual blocks, see below), as well as combinations of hyper-parameters, were evaluated to establish the choice that yields the highest classification accuracy on the validation set (corresponding to the lowest loss, measured in terms of binary cross-entropy).

The best performing network found has the following architecture: the two input layers (positional vectors and atom-types/B-values) are separately passed through a 1D convolution of thirty filters with linear activation functions (window size was set to 4) and subsequently concatenated to mix atom type and B-value-based patterns with the patterns learnt from the array of input vectors describing the ordered positional vectors; the data is then duplicated and passed into two collateral paths, the first one being a series of two residual blocks, each containing a series of two batch normalization layers with rectified linear unit activation functions followed by 1D convolutional layers, and the second being skip connections that join each residual block after passing its respective output data through a 1D convolutional layer. The paths are then joined together in a final residual block followed by a 1D convolution with a single sigmoid activation kernel and, lastly, the dimensionality of the model is collapsed in the final max pooling layer serving as the output for predicting whether or not a position is likely to correspond to a water molecule “hot-spot”. Altogether, the model comprises 34 141 trainable parameters (the model architecture is available for visual inspection in Supplementary figure 1, and the model summary is available in Supplementary file 2).

In order to minimize over-fitting, L2 regularization was applied (at the regularization rate λ=0.005) to the kernel parameter weights of the convolutional layers. Dropout was not found to improve the prediction accuracy and was not used. Additionally, in order to make the model more robust in the event of adversarial perturbations (or other non-standard inputs), the adversarial regularization wrapper from the TensorFlow Neural Structured Learning Framework was applied on top of the developed model (using the multiplier of the adversarial loss set to 0.2 & step size of 0.05) [45,46]. Lastly, in agreement with previous reports [47], we observed that the ‘adam’ optimizer did not guarantee a stable convergence of the loss function and it had to be replaced with the classic stochastic gradient descent (with a learning rate of 0.01).

### 2.4 Case examples for performance evaluation

In order to evaluate the performance of the model in a realistic setting, we sought examples of structures resolved using both high-resolution X-ray crystallography as compared to: 1) low-resolution X-ray crystallography; 2) NMR; and 3) Cryo-EM. Structures resolved using the latter techniques do not contain water molecules - they serve as the target structures for running predictions. The high-resolution structures, on the other hand, contain water molecule positions, which serve as a validation set for the predictions.

To this end, we arbitrarily selected “popular” proteins, for which many structures were available in PDB. We selected three high resolution (<1.8Å) structures resolved using X-ray crystallography: a carbonic anhydrase (resolved to 1.35Å; PDB accession 3M1K[48]), cyclophilin A (resolved to 1.63Å; PDB accession 2CPL [49]) and lysozyme (resolved to 1.4Å; PDB accession 2Q9D [50]). We used TopSearch [51] to search for the most structurally homologous (in terms of low root mean square deviation - RMSD) counterparts resolved to a lower resolution or not relying on X-ray diffraction. The resultant homologues found included: a crystal structure of a carbonic anhydrase resolved to 3.5Å (PDB accession 1G6V [52]; RMSD against 3M1K =0.73Å), an NMR structure of cyclophilin A (PDB accession 1oca [53]; RMSD against 2CPL=0.83Å) and a cryo-EM structure of a gamma-secretase assembly, which includes a lysozyme (resolved to 4.4Å; PDB accession 4UIS [54]; RMSD against 2Q9D =1.09Å). The rigid version of FATCAT [55] was used to superimpose structures on top of each other, so as to localize the binding sites of water molecules from the high-resolution structures within their low-resolution counterparts. Proteins exhibiting structural homology to the above listed structures (according to TopSearch) were excluded from the machine learning training set, so as to ensure independence of these test cases and avert the risk of reporting results obtained due to over-fitting.

### 2.5 Protein crystallization and structure determination

The N-terminal domain of *T. cruzi* PEX14 was expressed and purified as previously described [13]. Protein was crystallised at 40 mg/ml in crystallization buffer containing 0.2 M Na acetate, 0.1 M Tris.HCl at pH 8.5 with 30% (w/v) PEG 4000. Crystals were grown at 20 °C and cryoprotected in glycerol. The data was indexed using XDS [56] and scaled using XScale. The initial phases were obtained using molecular replacement carried out using Phaser [57], with *T. brucei* PEX-14 as the search model. Manual rebuilding using electron density maps was carried out in Coot [58]. Phenix Elbow [59] was used for obtaining restraints for small molecules in the crystallization conditions. Further refinements were carried out using Phenix Refine. 5 % of the reflections were used for cross-validation analysis. The final model will be deposited in the PDB; PDB code to be confirmed.. Figures were produced using Pymol [60], Coot, Affinity Designer and Inkscape.

## 3. Results & discussion

### 3.1 Characteristics of the generated negative class dataset

A comparison of water versus “non-water” (negative class samples) summary statistics is presented in Figure 2. The features examined included ‘distance to protein’s closest atom’ - the distance of the oxygen of the water molecule to the closest atom of the protein; ‘Mean temperature factor’ - mean temperature factor of all protein atoms in the neighborhood of the water molecule (<7.5Å away from the water molecule); ‘Maximum angle between protein’s atoms’ - the maximum angle between any pair of vectors pointing to the protein atoms interacting with the water molecule, serving as a measure of how deeply buried the position is within the protein’s solvation surface, and “number of interacting atoms” with the water molecule (number of atoms <7.5Å of the oxygen of the water molecule).

**Figure 2:**
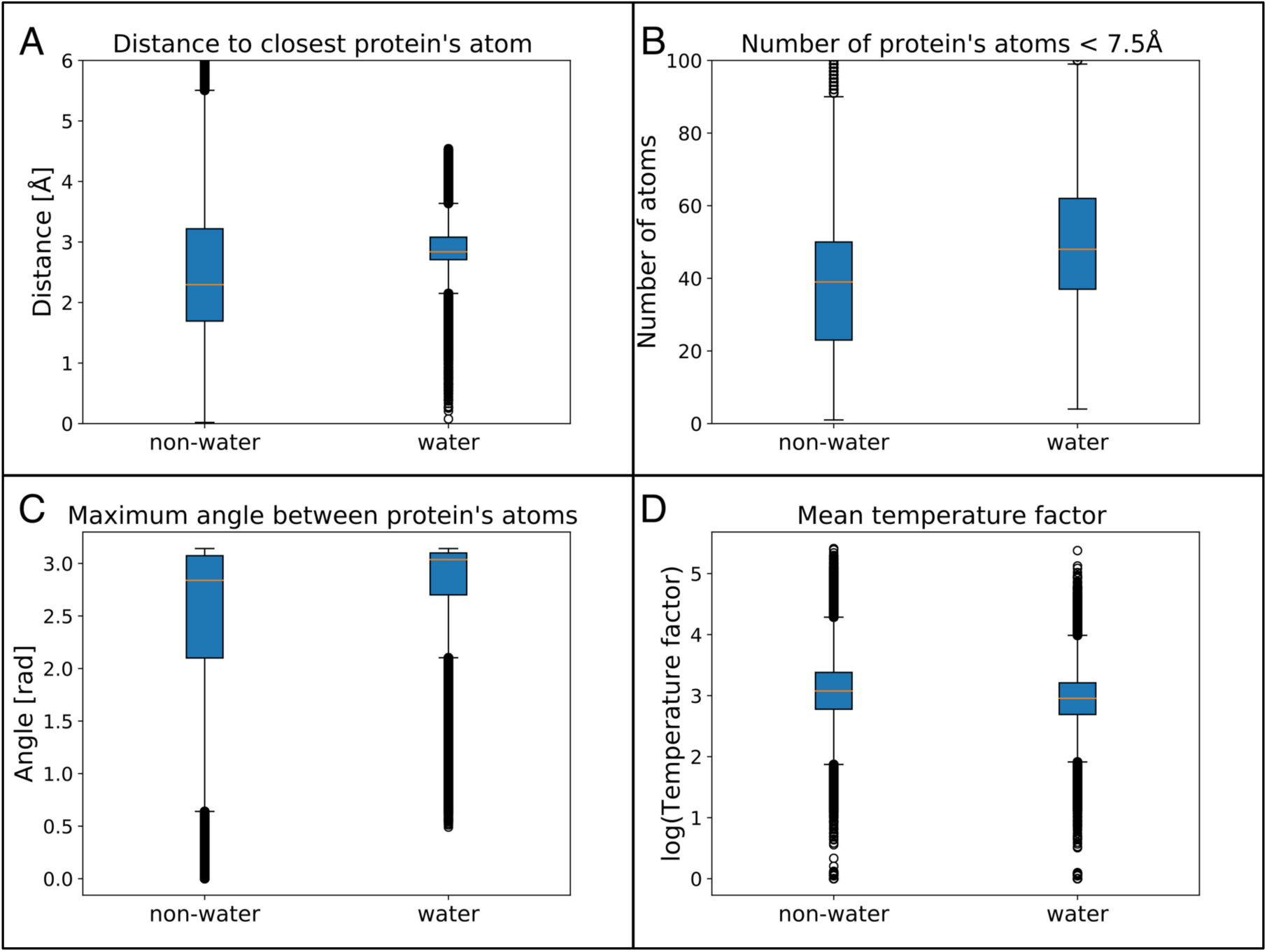
Features describing characteristics of positive (water molecules within PDB files) and negative (heuristically chosen non-water positions) class samples: A) distance to closest atom of the protein; B) Number of interacting protein’s atoms (<7.5Å); C) maximum angle between vectors pointing to any pair of protein’s atoms – as a measure of how deeply buried the position is; D) mean temperature factor of protein’s atoms in the neighborhood of the position (<7.5Å).

The majority of water molecules encoded within PDB files are between 2.6 and 3.2Å from the closest interacting atom of the protein. This range (1.8-3.3Å) is much broader for the points used as the negative class dataset but it extends in both directions (points closer and further away from the protein’s atoms) and at least 10% of negative samples lie exactly in the range most frequently occupied by water molecules. The mean temperature factor of protein’s atoms in the positions near water molecules is clearly lower than that of the negative class dataset but the overlap in the ranges of values measured for the two sets is over 75%. The maximum angle between any pair of vectors pointing to the protein’s atoms is generally higher (>2.7 rad) for water hot-spots but over 50% of positions used as the negative class samples are buried equally deep within the protein’s surface. Finally, we observed that water molecules typically interact (reside within 7.5Å of) with 35-60 protein atoms; roughly 40% of samples in the negative class dataset exhibit the same range.

Overall, while the negative class dataset contains many samples that appear to be trivially easy to predict as unlikely to be occupied by water molecules, a significant fraction still offers room for learning the non-obvious water hot-spot characteristics.

### 3.2 Performance of the final model

The network described in section 2.3 was trained for 100 epochs and achieved an accuracy of 94% on the test set (Figure 3A) and the area under the receiver operating characteristic curve of over 0.98 (Figure 3B). It is important to note that the model performance determined here applies only to the training dataset used in this study. The accuracy is expected to be highly sensitive to the number of “trivial” non-interacting sites the model was presented with during training. Likewise, filtering out the positive dataset to exclude water molecules for which there is insufficient evidence in the form of electron density clouds could further improve the classification accuracy. We have not performed this step since Nittinger et al. have shown that roughly 90% of water molecules encoded on the surfaces of proteins within PDB files were real [14]. While this fraction is already high, we have further increased it by excluding all water molecules not featuring interactions with at least two atoms of the protein structure (this was generally between 0-15% of water molecules encoded in each PDB file) or further away than 4.5Å of any atom of the protein. The main reason behind not using the dataset provided by Nittinger et al. is that it does not guarantee that the protein structures available are non-redundant – this could potentially lead to information leakage between folds of the cross validation and also the test set. Secondly, their dataset comprises only 2.3 million water molecules which is significantly smaller than the dataset we used.

**Figure 3:**
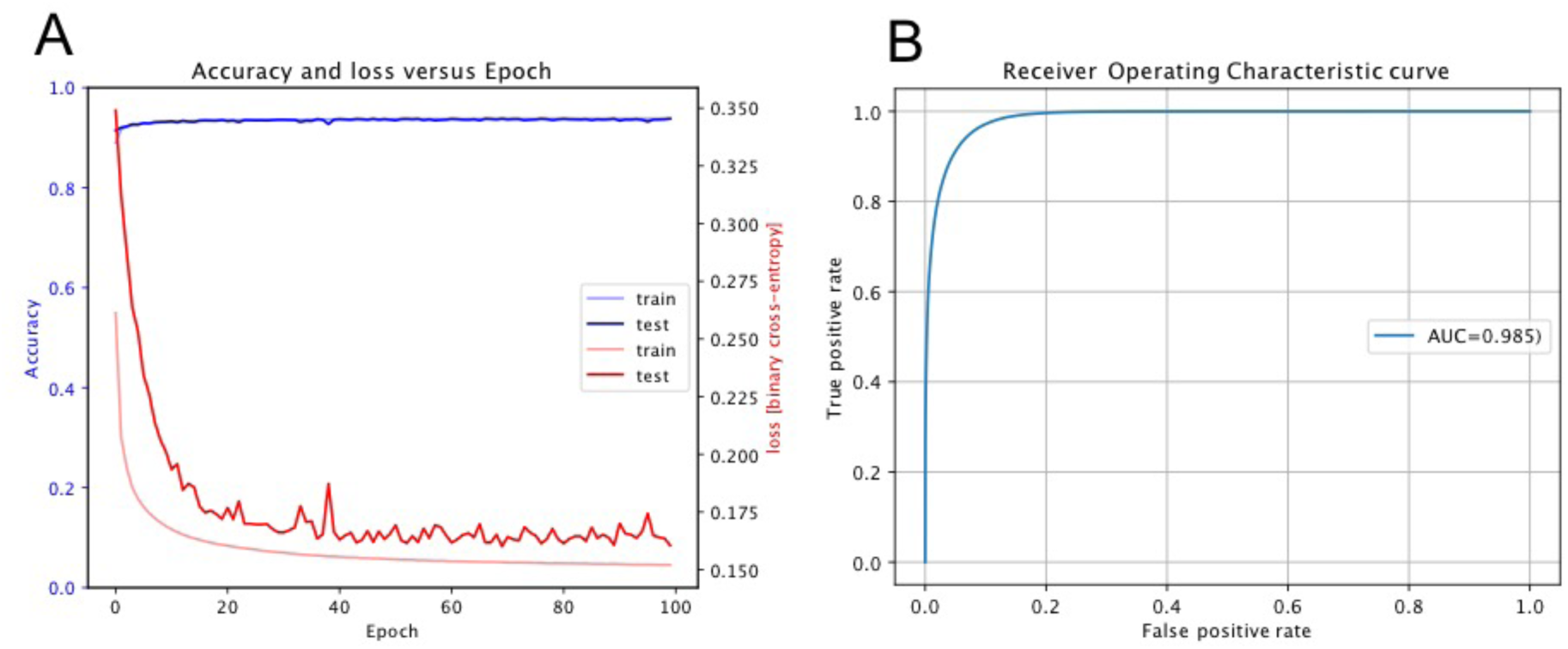
Performance of the final model. A) Left axis: model accuracy across training epochs (light blue and dark blue line for train and test set, respectively; Right axis: binary cross-entropy loss across training epochs (light red and dark red line for train and test set, respectively B) Receiver Operating Characteristic curve i.e. false-positive versus true positive rate; area under the curve= 0.985.

### 3.3 Characteristics of high-scoring water molecule positions

Using the prediction scores from the developed model, we examined the characteristics of the positions: unlikely to be interacting with waters (prediction score <0.5) or ever more likely to be water hot-spots (prediction scores 0.5-0.9, 0.9-0.99, 0.99-0.999, and >0.999). Unsurprisingly, we found that the high-scoring water hot-spots are buried within the protein’s surface (figure 4C) and surrounded densely by atoms of the protein (figure 4B); this increases the likelihood of creating multiple hydrogen bonds precisely at the configuration, which confers the most stable interaction. The normalized temperature factors also show a clear anti-correlation with prediction scores (figure 4D), however, based on the breadth of the distributions, this is not a major factor affecting the predictions. Interestingly, the highest scoring water hot-spots feature precisely the same distance to closest atoms as the average water molecule used in the training set (2.8Å; figure 4A). At this point, we think it is important to underline the fact that the features extracted here were not explicitly input into the model – they have been learnt from the raw data encoded as described in section 2.2.

**Figure 4:**
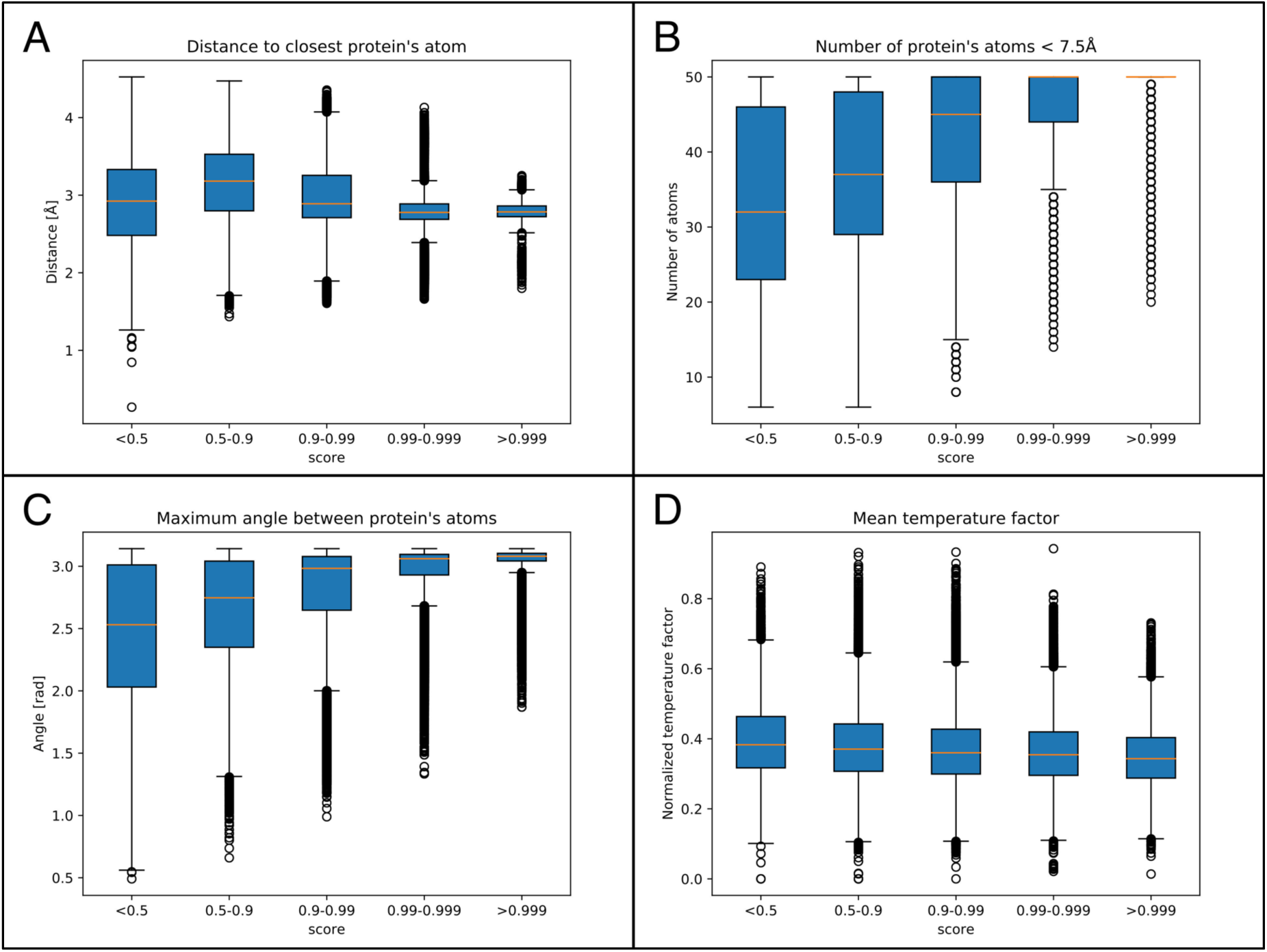
Features describing characteristics of water molecules according to the assigned hot spot prediction scores: A) distance to closest atom of the protein; B) Number of interacting protein atoms (<7.5Å); C) maximum angle between vectors pointing to any pair of protein atoms – as a measure of how deeply buried the position is; D) mean normalized temperature factor of protein atoms in the neighborhood of the position (<7.5Å).

### 3.4 Scanning structures without information on water to detect water hot-spots

In order to empirically validate the final model, we evaluated its performance by comparing the predicted water hot-spots within low-resolution X-ray, NMR and Cryo-EM structures against water molecules identified within structurally homologous high-resolution X-ray structures (table 1). Figure 5 depicts an example of one of the low-resolution structures, which was scanned for water hot-spots.

**Table 1:**
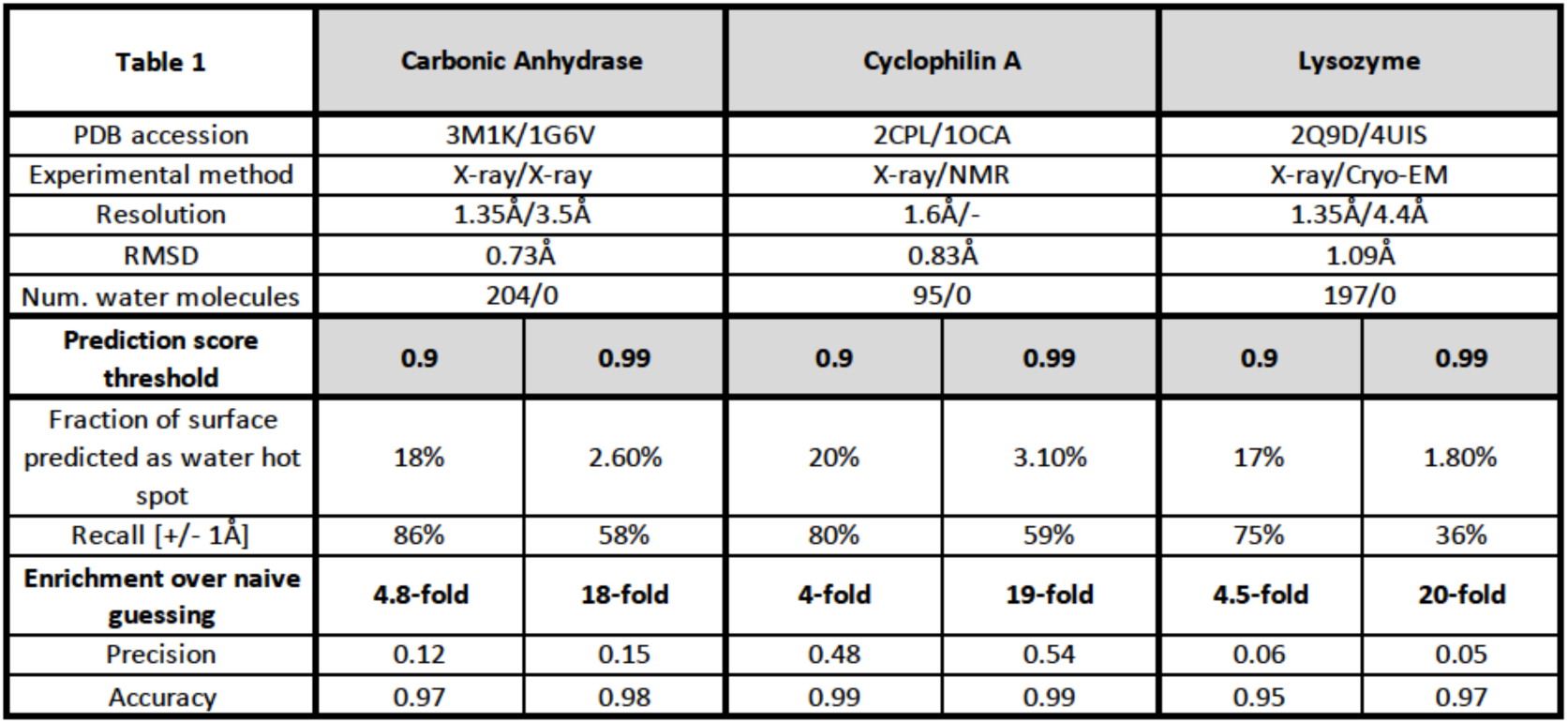
Model performance evaluation with empirical examples.

| Table 1 | Carbonic Anhydrase |  | Cyclophilin A |  | Lysozyme |  |
| --- | --- | --- | --- | --- | --- | --- |
| PDB accession | 3M1K/1G6V |  | 2CPL/1OCA |  | 2Q9D/4UIS |  |
| Experimental method | X-ray/X-ray |  | X-ray/NMR |  | X-ray/Cryo-EM |  |
| Resolution | 1.35Å/3.5Å |  | 1.6Å/- |  | 1.35Å/4.4Å |  |
| RMSD | 0.73Å |  | 0.83Å |  | 1.09Å |  |
| Num. water molecules | 204/0 |  | 95/0 |  | 197/0 |  |
| Prediction score threshold | 0.9 | 0.99 | 0.9 | 0.99 | 0.9 | 0.99 |
| Fraction of surface predicted as water hot spot | 18% | 2.60% | 20% | 3.10% | 17% | 1.80% |
| Recall [+/- 1Å] | 86% | 58% | 80% | 59% | 75% | 36% |
| Enrichment over naive guessing | 4.8-fold | 18-fold | 4-fold | 19-fold | 4.5-fold | 20-fold |
| Precision | 0.12 | 0.15 | 0.48 | 0.54 | 0.06 | 0.05 |
| Accuracy | 0.97 | 0.98 | 0.99 | 0.99 | 0.95 | 0.97 |

**Figure 5:**
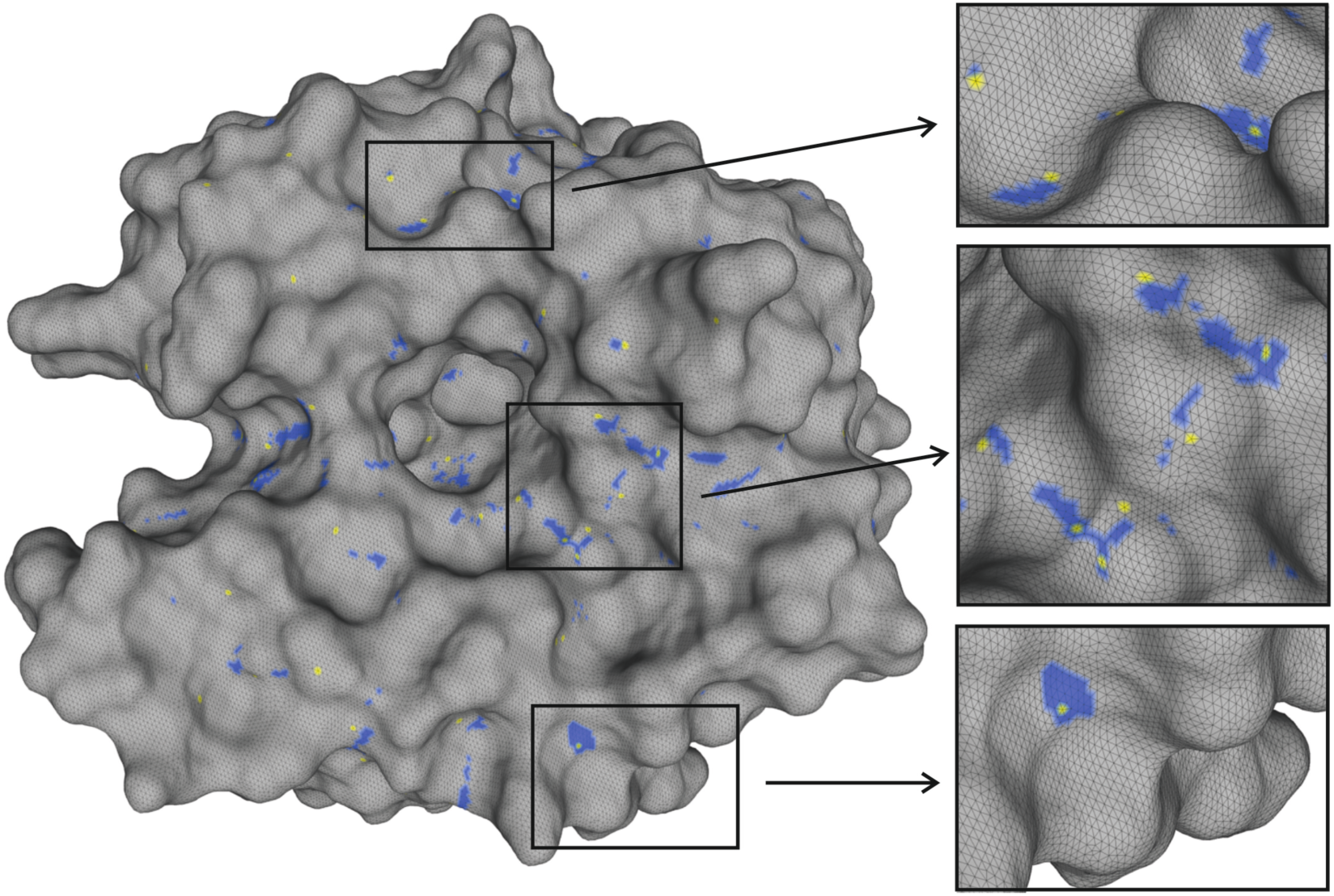
Solvation surface of low-resolution X-ray structure 1G6V scanned for water hot-spots (shaded blue); magenta points indicate positions of water molecules superimposed from the high-resolution structure 3M1K.

Scanning the solvation surface of an entire protein at sub-Angstrom resolution involves generating predictions for over 10 million points. This inevitably leads to over-predicting the number of hot-spot positions, however, selecting a strict prediction threshold for defining positive predictions can alleviate this problem. For example, when choosing the threshold to be 0.9, in the tested examples, the model predicts 17-20% of the protein’s surface to interact with water; this allows retrieving at least 75% of real water molecules encoded within the high-resolution X-ray structures, which already offers at least 4-fold enrichment over naive guessing (selecting positions on the protein surface at random). Increasing the threshold for positive predictions further (up to 0.99) reduces the number of false-positive predictions (2-3% of protein’s surface predicted to interact with waters) at the expense of a slightly reduced recall (this allowed retrieving 36-59% of water molecules encoded in the corresponding high-resolution X-ray structures); this offers an 18-20 -fold enrichment over naive guessing.

One explanation for some of the differences seen could be the intrinsic differences between the methods themselves. X-ray crystal structures are affected by many additional parameters, such as crystal packing, pH, temperature, and the presence of ions or other additives that may affect the presence of water molecules in the system. The presence of such additional molecules and molecular ions is currently not an input into our computational process, which analyzes the protein molecules present in a single unit cell of a solved structure; thus, the system could predict a water molecule in place of an ion or crystal contact, leading to a deviation from the expected results. The same is true for NMR structures, which are measured in aqueous solution; the conditions are very different to those *in silico* as well as to those in high-resolution X-ray structures, as used for the training set, due to different environments and higher intrinsic flexibility in solution. The flexibility seen in solution, along with the variety of conformations measurable in both NMR and Cryo-EM are not represented in the training set, which prevents the machine learning model from “seeing” and learning from alternative biologically relevant protein conformations. This inherent bias in protein conformations attributed to crystal structures is also likely to contribute to the disagreements between computational predictions obtained for non-crystallized proteins and the water positions identified in X-ray-based structures.

Further increasing the threshold for positive predictions to 0.999 yields in the order of several hundred positively predicted spots. These, according to the model, are the precise locations that feature the most stable interactions with water. Although many of these predicted sites are not corroborated by water molecules present in the corresponding high-resolution X-ray structures, in the following example we demonstrate that many of these are theoretically good binding sites for water and in fact, in some cases corroborated by electron density clouds.

### 3.5 Application of the method to a newly crystal resolved structure

A further, previously unpublished, drug discovery target structure of the N-terminal domain of *T. cruzi* PEX14 was also critically compared to the results as determined using the final model. Taking the highest scoring predictions (score >0.999), out of the 222 water molecules considered, 60 were considered to have been predicted correctly meaning that 27% of real water molecules present within the defined grid were captured, giving a 150-fold enrichment over naive guessing. With a score threshold of >0.99, 70% of the water molecules are captured at the cost, as before, of a higher false-positive rate: 2.6% of the surface is covered by positive predictions, giving a 27-fold enrichment over naive guessing. These results are consistent with what we would expect from the model, giving better results than those above, where water positions in different, albeit similar, structures were compared. Manual inspection of the results, however, highlighted some differences between calculated and experimental as could be expected, and one further, as yet unaccounted for, difference, that could affect the apparent statistics of the model.

Figure 6 A and B show example comparisons of the experimentally determined waters and the predicted water positions according to the hotWater model. In A, the calculated and experimental match very well, whilst in B, both false negatives and false positives are illustrated. It is worth pointing out that the false negative positions did not meet the stringent >0.999 score threshold, which suggests that these sites feature a relatively lower affinity towards water than the other positively captured sites. As previously mentioned, crystallography often includes the addition of additives and cryo-protectants which could affect the apparent accuracy of the model presented. Figure 6 C illustrates this case; the algorithm indicates a high probability of water being present, as shown by many water positions being modelled. However, a glycerol OH is present in the crystal structure instead, leading these water positions to be considered as false positives. Were there to be no glycerol present in the condition, it is plausible that a water molecule would indeed take the place of the –OH group.

**Figure 6:**
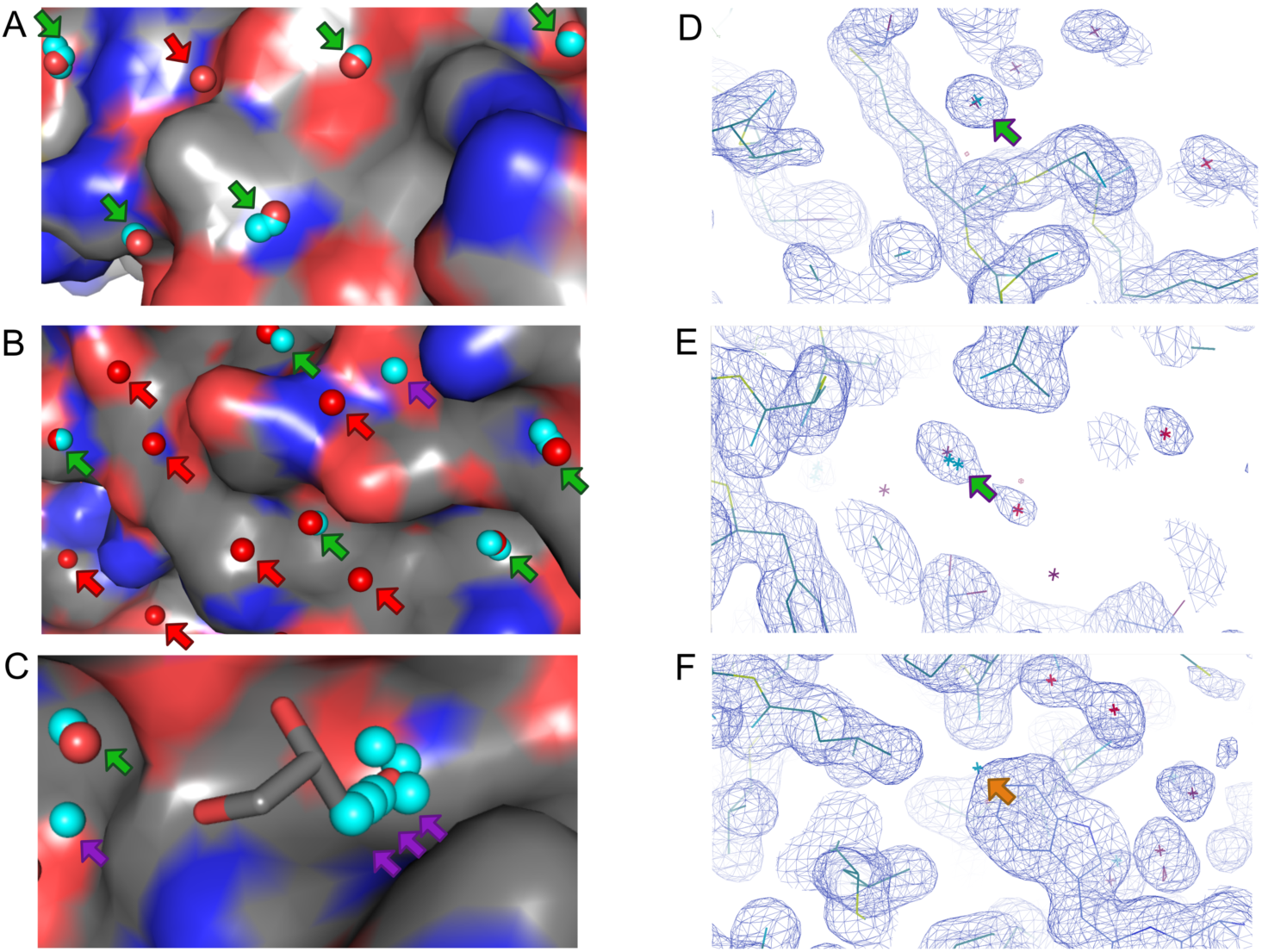
the hotWater model applied to a new crystal structure of N-terminal domain of PEX14. Green arrows indicate true positive predictions, red arrows indicate false negative predictions, purple arrows indicate false positive predictions and the orange arrow indicates a special case: A) area of structure covered primarily by true positive predictions; B) Area of structure with false negative and false positive predictions; C) site predicted to harbor many water molecules which are instead occupied by a glycerol OH in the real crystal structure and thus deemed as false positive; D-E) Water molecules predicted using the hotWater model, that are not present in the unit cell (thus showing as false positives) but are present in the neighboring unit cell. These are therefore ‘false false positives’. F) a water is predicted using the algorithm, but in the neighboring unit cell, there is instead the tryptophan present, preventing the presence of a water. In D-F) Unit cell waters shown in red, neighboring unit cell waters in purple and predicted waters are shown in turquoise.

The other factor that would affect the statistics of the model is illustrated in Figure 6 D-F. In crystallography, the structure of a unit cell is calculated within the electron density map, which is then iteratively used in the refinement calculations and further manually refined to repeatedly improve the structure. The unit cell motif repeats periodically throughout the crystal lattice, with the edge of one unit cell touching the edge of the next unit cell in a tessellating pattern. When only one unit cell is inspected, the molecules at the edge of the neighboring unit cell are therefore disregarded. In Figure 6 D-F, three cases of this are shown pictorially. The unit cell water molecules provided are shown in bright red, whilst those of the neighboring unit cell are shown in a deeper purple. All three cases show purported false positives; that is, a water molecule is predicted but none is seen in the PDB of the unit cell. However, when we manually inspect the crystal structure, so that neighboring unit cells’ electron densities can also be inspected carefully, we see that in the cases of D and E, there is indeed a water molecule where the model predicted. In F, there is instead an alternative molecule seen in the unit cell, the aromatic group of a tryptophan sidechain that would clash with the predicted water. This means that whilst it is possible that a water could sit there, it would never be seen in the crystal structure due to the crystal packing. As ‘false false positives’ like these (Figure 6 C-F) are included in the calculation of the statistics as false positives, the model’s apparent statistics indicate a higher false positive rate than is truly the case.

Another use in the case of a new structure is as a tool during refinement. We input the backbone manual refinement with no waters present into the deep learning algorithm and used the output in a refinement calculation using Phenix. This gave an Rfree value (used to measure the fit of a model to the electron density, where a lower Rfree value represents a better fit) of 0.2318. In comparison, a manual refinement before the addition of waters had an Rfree of 0.2416, larger by around 1 %. This is at a resolution of 1.6 Å. We then compared the algorithm with the Coot Find Waters tool and the Phenix Update Waters option in the refinement process at a variety of resolution cut-offs (Figure 7). It was found that at cut-offs at lower resolution (>2.6Å), no waters were found using Phenix and a maximum of 6 water molecules using Coot. The HotWater algorithm has a recall of 0.27 at all resolutions where the protein structure can be fitted as it relies purely on the PDB. This recall is only matched by Phenix at a resolution of 2.5Å and was not matched by the Coot Find Waters (with default settings) at any resolutions tested. The precision of the hotWater algorithm is, however, lower than that for the two algorithms that use electron density. This is of course understandable given that electron density is fixed evidence, whilst our algorithm is extrapolating from other datasets. It is conceivable however, that with implementation of electron density data into the training set, our algorithm would have reduced false positives due to the unit cell boundary and have even better recall and precision statistics. This would be a valid approach for further improvement of the software for the crystallography community.

**Figure 7:**
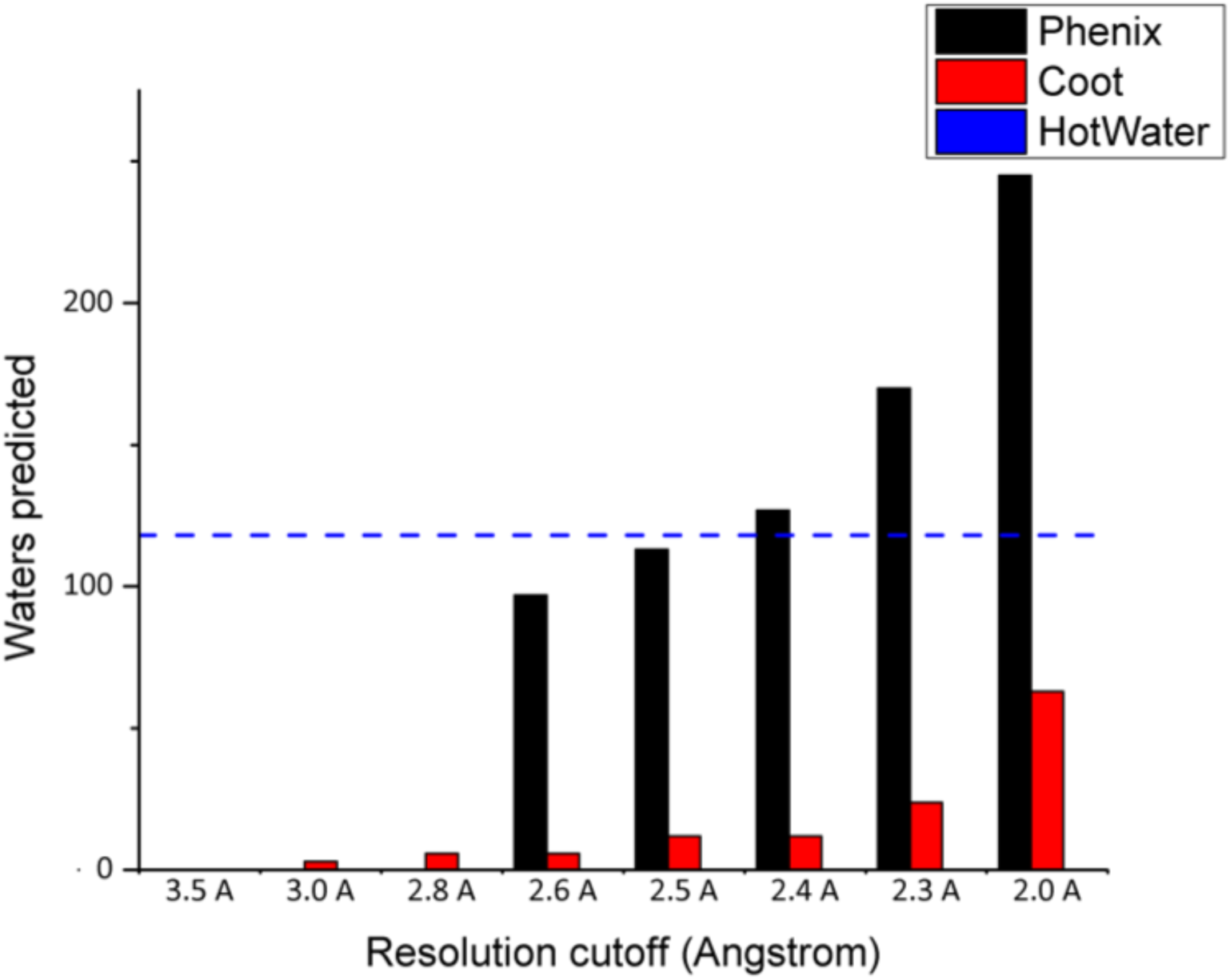
Depiction of waters predicted by different algorithms at a variety of resolution cut-offs. The hotWater algorithm is not affected by resolution of the electron density map, provided that the protein backbone can be fitted and, as such, is shown as a dashed blue line.

### 3.6 Direct applications for drug discovery

For use in drug discovery, waters that are particularly favorable are sometimes deemed to be ‘water hot-spots’ or ‘magic water’ molecules. This means that they are so fundamental in terms of energetic considerations that drug discovery attempts would be ill-advised to consider displacing them. They are often found in or at the edge of active site regions and, instead of being displaced, are normally considered by drug discovery teams as part of the protein surface, meaning that ligands are designed to maintain the position of that water and maximize interactions with it. Our algorithm allows the ranking of water molecules considering the strength and types of interactions, based on how consistently similar sites are occupied by water molecules in the training set. By using a particularly high or narrow threshold, only the most dominant waters would be highlighted, allowing scientists to analyze the presence of potential ‘water hot-spots’ within their structures. Currently the standard way to determine their presence is to solve several crystal structures, thus highlighting their repeated presence, or by synthesizing molecule that should extend into the area of the water to displace it and finding a reduced activity, despite a significant increase being expected. This is costly in both time and financial terms and a computational solution to this would be of great benefit to the drug discovery community.

Another trend that we have seen in the analysis of the structural outputs of the hotWater algorithm was the co-localization of predicted waters with oxygen atoms of small molecules or ligands in the X-ray structures. In comparisons with both of the recently solved structures, this was seen. In the tcPEX14 structure, a water was placed in the position of the oxygen atom of glycerol, whilst in the IMP-13 structure (PDB: 6RZR [61]), waters were predicted in the locations of the oxygen atoms of the substrate. Based on this, studying the predicted water positions in and near interesting binding sites could really enrich the information possessed by structural biologists and aid in design of molecules that maximize the interactions of the predicted water molecules.

### 3.7 Application of the method to urgent targets with structures solved by other methods

In recent months, the news has been dominated by the novel coronavirus, SARS-CoV-2, and the disease caused by it, COVID-19. The high rate of infection and severe health problems associated have led to vast changes to the way the world functions on a social, cultural and medical level. Scientists have rapidly solved the structure of and RNA polymerase in complex with its cofactors (PDB: 7BTF [62]) using cryo-EM at 2.9Å resolution. Application of the hotWater algorithm to this structure highlighted potential sites of key water molecules that could be exploited in drug discovery processes. This solvated structure could give scientists a better understanding of the solvation sphere of the protein and how this could affect its behavior and binding.

## Conclusions

Water binding site information is of critical importance in understanding the mechanisms of the protein’s action and in drug design. Current in silico fragment-based screening relies solely on computationally demanding, whilst still inaccurate physical force field models [63]. We have developed a machine-learning model for determining the water hot spots on a protein’s surface, that has been shown to work on previously published and new structures. This will help to determine water hotspots that are often critical to structure-based drug development. We envisage that our approach can also be extended to other small molecules and provide a novel deep learning tool for in silico screening and drug design. Our model can also serve to quickly, ab-initio solvate crystal structures during refinement even before the electron density is searched for water densities. Thus, it allows the elimination of a fraction of model bias typical in crystallographic model building.

## Acknowledgements

We’d like to thank the Frishman, Popowicz and Sattler groups for their support and helpful discussion. We are grateful to the Protein Expression and Purification Facility and the X-ray Crystallography Platform at Helmholtz Zentrum München for support. C.A.S., G.M.P. and M.S. acknowledge funding from the European Union’s Framework Programme for Research and Innovation Horizon 2020 (2014-2020) under the Marie Skłodowska-Curie Grant Agreement No. 675555, Accelerated Early staGe drug discovery (AEGIS).

## Supplementary figure

**Supplementary Figure 1:**
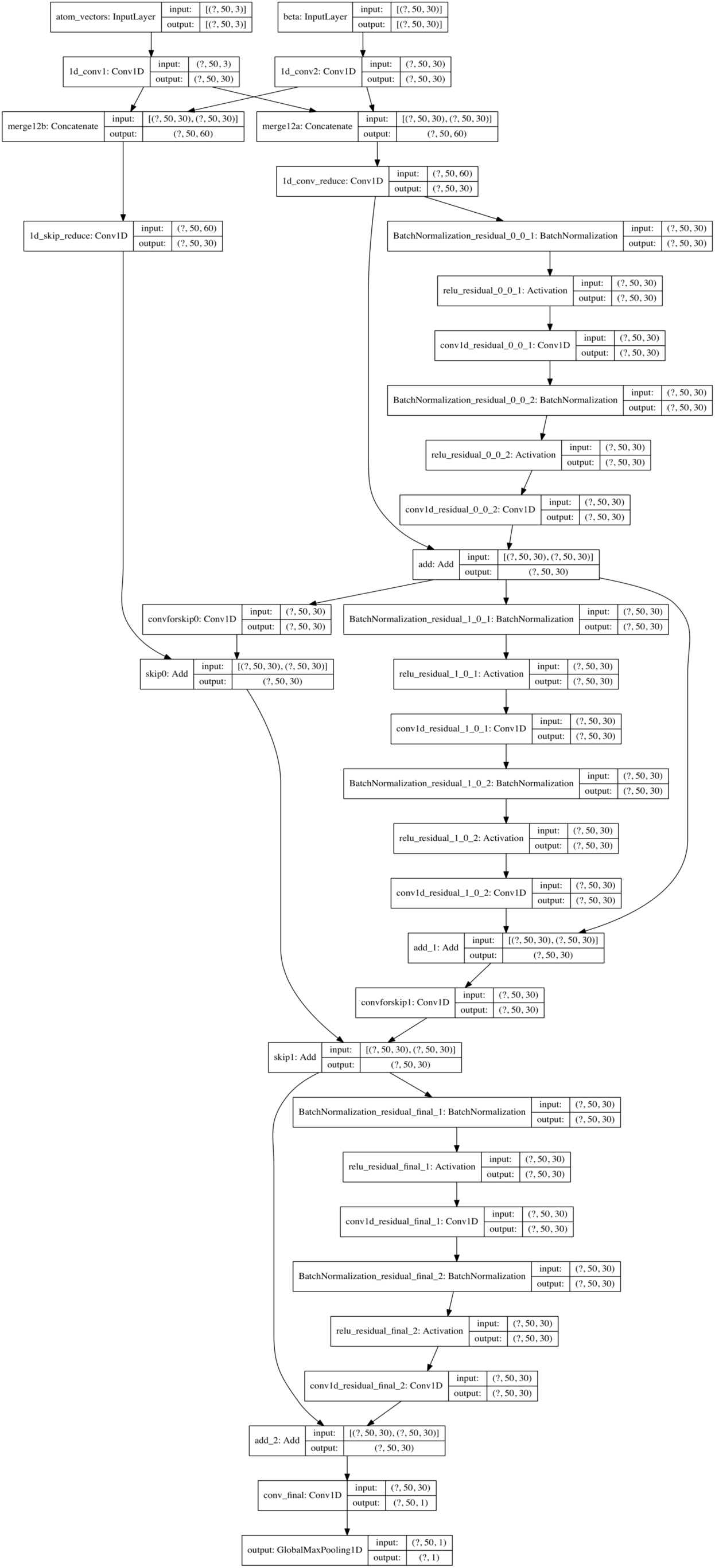
Deep residual neural network model summary.

